# Urban non-point source pollutants cause microbial community homogenization via increasing deterministic processes

**DOI:** 10.1101/2023.08.21.553702

**Authors:** Haizhou Li, Xiangyu Fan, Zhiwei He, Jing Fu, Yuekai Wang, Jinchen Yu, Shanshan Yang, Jiawang Wu, Li Wu, Jin Zhou

## Abstract

Urbanization significantly impacts the community structure of aquatic organisms and poses a major threat to river biodiversity. However, the extent to which urbanization is linked to the homogenization of microbial communities and the underlying mechanisms remains poorly understood. In this study, we investigated the bacterial and archaeal communities from cities and neighboring natural rivers across river network located in the Qinling Mountains, Northwest China, and further investigated the alpha and beta diversity patterns and the mechanisms influenced by urbanization. We found that the influx of urban non-point source pollutants created a eutrophic condition, and enhanced the urban river microbial populations. Meanwhile, the rapid urbanization tends to decrease the overall habitat heterogeneity, and imposed stronger homogeneous selection and caused microbial communities biotic homogenization. The mechanisms of biotic homogenization can be attributed to modulating generalist/specialist species and invasion of nonnative species. For instance, the urban river had a greater proportion of fast-growing bacteria, algae, nitrifiers, PAH-degrading bacteria, pathogens, fecal bacteria and antibiotic-resistant bacteria than natural river ecosystems. Overall, urbanization leads to a more uniform river biosphere, causing the extinction of unique local species and a subsequent decrease in the regional species pool.

## 1. Introduction

Rivers have been a preferred location for human settlements throughout history (1). However, in recent decades, human activities such as rapid urbanization and industrialization have discharged multiple anthropogenic contaminants and non-native (exotic) microorganisms into the river (2, 3). These contaminants can be either organic (e.g., antibiotics, polycyclic aromatic hydrocarbons, fertilizers, pesticides, manure) (4-7) or inorganic contaminants (e.g., nitrate, phosphorus, heavy metal) (8-10). According to the United Nations, 68% of the global population will live in cities by 2050 (11). Accelerated urbanization and intense human activities substantially influence the community composition and function of the river microbiome, and posing a considerable risk to riverine ecosystems. For example, centralized wastewater treatment plants (WWTPs) represent an important and common urban river pollution source. Evidences showed that effluents from WWTPs exert adverse effects on the functionality of river ecosystems, often leading to the deterioration of water quality, habitat degradation, and a reduction in the richness and diversity of benthic and sedimentary microbial communities (12-15).

WWTPs are a common source of point-source pollution in the river ecosystem and have become a global aquatic environmental issue. However, the immense industrial and domestic sewage are considered as the primary contributors to non-point source pollution (2, 16). Point-source pollution is easy to identify, but non-point source pollution is harder to identify and harder to address. The United States Environmental Protection Agency indicates non-point source pollution remains the primary unresolved issue affecting river water quality, yet there are little data on the impacts of widespread non-point source pollution on microbial communities in river ecosystems (17, 18). The potential ecological and health influences of non-point source pollution in river ecosystems require further research and monitoring.

Here, we focus on a large-scale river network, which flows out of the Qinling Mountains, where it eventually empties into the Yangtze River (Fig. 1). The Qinling Mountains, located in Northwest China, serve as a geographically important north-south boundary and natural barrier for the eastward flow of moisture from the Tibetan Plateau, providing a significant source of freshwater for downstream human communities. The distinctive feature of the river network is that it passes numerous cities (a current population of approximately 11.06 million) (19) and national nature reserves. Establishing nature reserves is an effective measure to avoid human activity disturbances (20, 21). This river network provides a useful model for understanding the impact of non-point source pollution on the river ecosystem.

**Figure 1.**
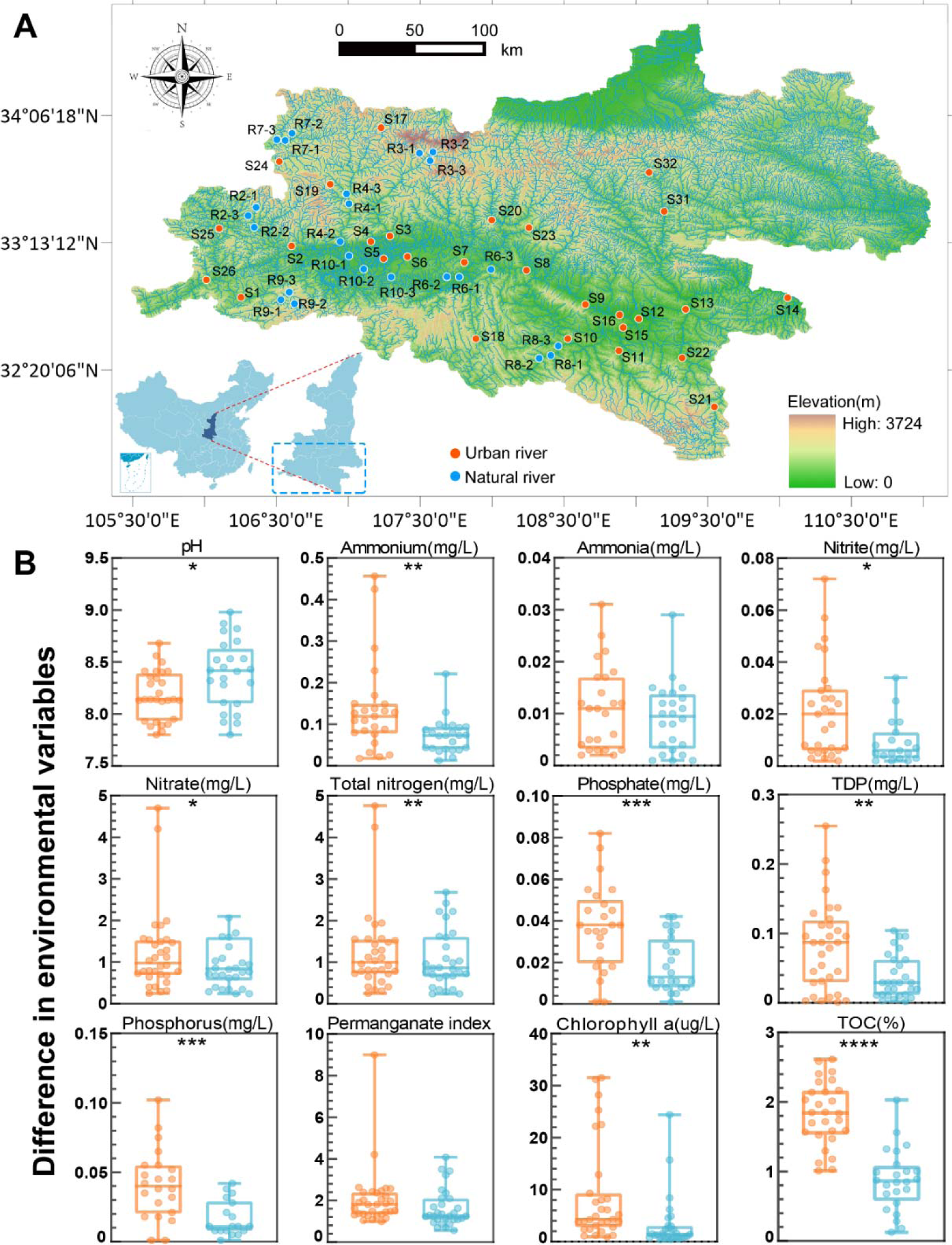
(A) Location of the collected river network sediment samples. The Inset map shows the extent of the study area relative to China. (B) Differences in environmental variables between the urban and natural river areas. Orange circles indicate the urban river sampling sites, and blue circles indicate the natural river sampling sites, respectively.

In this study, the collected river surface sediment samples (0–2 cm deep) were in eutrophic waterbodies (highly urbanized rivers) and oligotrophic or mesotrophic waterbodies (undisturbed nature reserve rivers), based on trophic state index (TSI) (Fig. 1, Dataset 1B). River sediments are considered as the major habitat of microbes in the aquatic environment (22, 23). Therefore, sediment-associated microorganisms drive essential ecological processes and contribute significantly to river function (24). Here, these collected river surface sediments represent diverse non-point source pollution gradients, and creating multiple niches in this river network. We obtained millions of 16S rRNA tags sequences from these sediments by high-throughput sequencing, aiming to (i) compare the community composition, diversity and function of bacteria and archaea in urban river with natural river ecosystems; (ii) evaluate the environmental factors linked to the river microbiome; (iii) identify the river microbial taxa (bacteria, archaea) community assembly processes; and (iv) assess the microbial functional attributes characterizing in river ecosystems associated with urbanization, eutrophication, antibiotic resistance, fecal contamination, pathogenesis, nutrient cycling, and abiotic stress.

## 2. Materials and methods

### 2.1 Study area, sampling procedure, and physicochemical analysis

Sampling was conducted in August-September 2022. From each sampling site on three sampling occasions, water and sediment samples were collected within approximately 2 square meters at each site. The geographical data (longitude, latitude, and distance between sampling sites) were recorded by a GPS device (Dataset 1A). The water quality parameters, including water temperature, pH, dissolved oxygen (DO), water clarity of Secchi depth, transparency (SD), turbidity (TUB), total nitrogen (TN), nitrate-nitrogen (NO ^-^), ammonium-nitrogen (NH ^+^), total phosphors (TP), total dissolved phosphorus (TDP), phosphate, permanganate index, volatile phenol, *Chlorophyll*-a concentration were measured according to Environmental Quality Standards for Surface Water (Ministry of Ecology and Environment of People’s Republic of China, GB3838-2002). Sediment total organic carbon (TOC) was measured by the multi N/C 3100 (Analytik Jena). Each sample was performed in triplicate, and the values were expressed as the mean (n = 3). To evaluate the trophic status of each site, we calculated a trophic state index (TSI) based on the concentrations of TN, TP, Chl-a and SD in the water environment (25, 26). Five trophic levels were defined: oligotrophic (TSI < 30), mesotrophic (30 ≤ TSI ≤50), slightly eutrophic (50 < TSI ≤ 60), moderately eutrophic (60 < TSI ≤70) and highly eutrophic (TSI >70) (25). For detailed information on the environmental variables of the trophic groups, see Dataset 1B.

### 2.2 Sediment cell counts

Catalyzed reporter deposition-fluorescent *in situ* hybridization (CARD-FISH) was used to quantify the cell densities based on our previous studies (27). The probe-labeling peroxidases were EUB338 (bacteria, 5’-GCW GCC WCC CGT AGG WGT-3’) and Arch915 (archaea, 5’-GTG CTC CCC CGC CAA TTC CT-3’). Probe NON338 (5’-ACT CCT ACG GGA GGC AGC-3’) was used as a control. Frozen samples (0.5 g) were fixed with 4% paraformaldehyde for 24 h at room temperature. The fixed sediments were washed three times by centrifugation (8,000 × *g* for 10 min) using phosphate buffered saline (PBS) at 4 °C and stored in an ethanol/PBS buffer (1:1) at -20 °C for further processing. Later, 100 μL of fixed sediment was diluted with 900 μL ethanol/PBS buffer (1:1) and dispersed using ultrasound. Then, 20 μL of the dispersed sediment was further diluted in 20 mL of Milli-Q filtered water. The suspended sediments were filtered onto polycarbonate filters, and 0.1% low melting point agarose was dripped onto the filters and dried at 46 °C in an incubator. The microbes were permeabilized using 15 μg/mL proteinase K. Then, 3% H_2_O_2_ was used to inactivate the endogenous peroxidases.

For hybridization, filters were placed in a tube and mixed with 500 μL hybridization solution (10% dextran sulfate, 2% blocking reagent (Roche, Germany), 0.1% (w/v) sodium dodecyl sulfate, 20 mM Tris–HCl [pH 8.0], 0.9 M NaCl and formamide) and 1 μL of probe working solution (final concentration, 0.028 μM) (28). Microorganisms were hybridized for at least 60 min on a rotor at 46 °C, and then the filters were washed twice using washing solution (28) (0.01% SDS, 5 mM EDTA [pH 8.0], 20 mM Tris–HCl [pH 8.0] and 3 mM NaCl) at 48 °C for 20 min. After washing, filters were mixed with 1,000 μL of amplification solution (0.0015% H_2_O_2_, 1×PBS [pH 7.4], 0.1% (w/v) blocking reagent) and 1 μL of Alexa 488-labelled tyramides (Life Technologies^TM^, Thermo Fisher, USA). The probes were incubated at 46 °C in amplification solution for at least 30 min in the dark.

For second hybridizations, the first probe-labelling peroxidase was inactivated by incubating the filter sections in 0.01 M HCl for 10 min at room temperature and washing the sections with 50 mL of Milli-Q water. Then, the CARD-FISH protocol was repeated two times with the same filter sections by using different probes. The second hybridization was performed using Alexa 647-labelled tyramides (Life Technologies, Thermo Fisher, USA). Finally, all microorganisms were stained using DAPI and mounted with ProLong Gold Antifade reagent (Life Technologies, Carlsbad, California, USA). Cell counting was performed using ImageJ.

### 2.3 High-throughput sequencing

Frozen sample was aseptically subsampled for DNA extraction. In addition, a negative control (5 mL of DNase water) was included with each batch of subsamples and subjected to DNA extraction. DNA was extracted from each subsample using the E.Z.N.A.® soil DNA kit (Omega Bio-tek, Norcross, Georgia, USA), according to the manufacturer’s instructions. The final DNA concentration was determined by a NanoDrop 2000 (Thermo Scientific, Waltham, USA), and DNA quality was checked by 1% agarose gel electrophoresis. The sequencing steps were conducted by Majorbio Bio-Pharm Technology Co., Ltd. (Shanghai, China). The bacterial/archaeal universal primers were 515F and 806R (29, 30), targeting the 16S rRNA gene V4 region. The polymerase chain reaction (PCR) protocol was performed according to previously described methods (31). The PCR products were extracted from a 2% agarose gel and further purified using an AxyPrep DNA Gel Extraction Kit (Axygen Biosciences, USA) and quantified using Quanti Fluor™-ST (Promega, USA). Purified amplicons were pooled in equimolar amounts and paired-end sequenced (2 × 300) on an Illumina MiSeq platform (Illumina, San Diego, California, USA). Raw fastq files were demultiplexed, quality filtered by Trimmomatic and merged by FLASH. OTUs were clustered with a 97% similarity cut-off using UPARSE, and chimeric sequences were identified and removed using UCHIME. To remove the effects of sampling intensity (number of reads) on comparisons of richness estimates, we randomly subsampled a fixed number of reads (n= 10, 000) for each sample. Taxonomic assignment was performed using the SILVA 16S rRNA gene database (version 138). The distance-based maximum likelihood was used for phylogenetic analysis. Bootstrap analysis was performed using 1,000 replications. The Mega software had been used for computing phylogenetic trees. Alpha-diversity metrics (observed OTU richness, Chao1 richness, Shannon and Simpson diversity indices, and equitability index) were calculated using Mothur (32).

Here, the “niche breadth” approach (33) was used to quantify habitat specialization. The formula is described below:

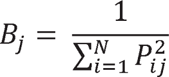

where *B_j_* is the habitat niche breadth of OTUj in a metacommunity; and P_ij_ is the proportion of OTUj in a given site i, and N is the total number of sites. A higher B value of OTUs indicates a wider habitat niche breadth. In contrast, lower *B* values indicate that the OTUs occurred in fewer habitats and were unevenly distributed (34). The average of B values from all OTUs in one community was regarded as the community-level B-value (*Bcom*) (35). We calculated the *B* value based on species abundance and evenness for NHF and SCS sediment samples (36).

Furthermore, in order to identify microbial ecological functions at the studied sites, the obtained OTUs were compared with the PICRUSt2 (37, 38) and FAPROTAX database (39, 40) to predict metabolic functions of the microbial community.

### 2.4 Screening for antibiotic resistant bacteria from river sediments

The approach that has been used to evaluate the status of antibiotic resistance in the environment involves the calculation of resistance percentage, which is based on the ratio between the number of bacteria that are able to grow on culture media that are supplemented with antibiotics at doses close to the minimal inhibitory concentration (MIC) and the number of bacteria that grow on antibiotic-free media (41, 42). The approach of heterotrophic plate count (HPC) method (43) was used to determine the concentration of antibiotic resistant bacteria in the collected river surface sediments. 0.1 g sediment samples were dissolved in 10 mL of 0.85% sterilized normal saline, and diluted 10 × fold for immediate processing. 100 μL diluted samples were cultured onto plate count agar medium. Medium contained four-class plates, 50.4 μg/mL sulfamethoxazole, 16 μg/mL tetracycline, 15 μg/mL azithromycin and no antibiotics. The media was added antifungal agent 200 μg/mL cycloheximide according to Gao el al., (2012) and Brooks et al. (2007) method. Total heterotrophic cultivable bacterial population was determined by plate count agar medium without antibiotics. The selected concentrations of antibiotics on the media were determined according to the criteria of the Clinical and Laboratory Standards Institute (CLSI) guidelines (2015) and the previous study (43-45). Plates were incubated for 48–72 h at 37 °C (43, 46). Duplicate counting was performed, and colony-forming units (CFU) were calculated.

### 2.5 Real-time quantitative PCR (qPCR)

Absolute quantification standard curve method was used to quantify the antibiotic resistant genes in the river sediment samples based on our previous studies (47). The real-time quantitative PCR (q-PCR) (ABI7500, USA) using SYBR 2×RealUniversal PreMix (Tiangen Biotech). qPCR was performed to assess the abundance of the macrolide, tetracycline and sulfonamide antibiotic resistant genes (ARGs). The sequences of the primers and the reaction conditions used in this study are presented in Table S1 in the Supporting Information (SI) section.

The PCR reactions were performed in a 20-μL reaction mixture that comprised 10 μL of SYBR RealUniversal PreMix (2 ×), 0.4 μL of each type of primer (10/20 μM), and 1 μL of a DNA template (5 ng). The thermocycling protocol consisted of an initial denaturation for 1 min at 95 °C, followed by 40 cycles of denaturation (95 °C, 10 s) and annealing at the temperatures in Tables S1. The specificity of the PCR products was checked by a melting curve analysis. All qPCR assays were performed in triplicate for each gene,

The q-PCR standards were synthesized by cloning target genes into *Escherichia coli* DH5a using a PMD18-T vector Cloning Kit (TaKaRa Biotechnology, China). Vector concentrations were measured using NanoDrop, and the quantities of target genes per μL of solution were obtained basing on the length of plasmid and target gene sequence. Six-point calibration curves were generated by 10-fold serial dilutions of the plasmid carrying ARGs. All PCR reactions were run in parallel with serially diluted q-PCR standards and DNA-free water as the negative controls.

### 2.6 Statistical analysis

Null model analysis was carried out using the framework described by Stegen et al. (2012, 2013) to classify community pairs into underlying drivers of species sorting (or selection), dispersal limitation, homogeneous dispersal, and drift. The variation of both phylogenetic diversity and taxonomic diversity was measured using null model-based phylogenetic and taxonomic β-diversity metrics, namely β-nearest taxon index (βNTI) and Bray–Curtis-based Raup–Crick (RC_Bray_). A significant deviation (i.e., |βNTI| > 2) indicates the dominance of selection processes. βNTI < −2 indicates significantly less phylogenetic turnover than expected (i.e., homogeneous selection) while βNTI > +2 indicates significantly more phylogenetic turnover than expected (i.e., variable selection). Subsequently, RC_Bray_ was used to further partition the pairwise comparisons that were not assigned to selection (i.e., |βNTI| < 2). The relative influence of homogenizing dispersal was quantified as the fraction of pairwise comparisons with |βNTI| < 2 and RC_Bray_ < –0.95. Dispersal limitation was quantified as the fraction of pairwise comparisons with |βNTI| < 2 and RC_Bray_ > 0.95. The fractions of all pairwise comparisons with |βNTI| < 2 and |RC_Bray_| < 0.95 were used to estimate influence of “undominated” assembly, which mostly consists of weak selection, weak dispersal, diversification, and/or drift.

A neutral community model was used to determine the contribution of stochastic processes to microbial community assembly by predicting the relationship between the frequency with which taxa occur in a set of local communities and their abundance across the wider metacommunity (48). The model predicts that abundant taxa are more likely to be dispersed by chance and widespread across metacommunity, while rare taxa would be lost in different local communities due to ecological drift. In the model, the estimated migration rate is a parameter for evaluating the probability that a random loss of an individual in a local community would be replaced by dispersal from the metacommunity, and, therefore, is a measure of dispersal limitation. Higher *m* values indicate that microbial communities are less dispersal limited. The formula is as follows:

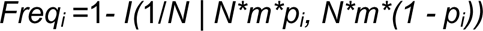

where *Freq_i_*is the occurrence frequency of taxon *i* across communities; *N* is the number of individuals per community; *m* is the estimated migration rate; *p_i_*is the average relative abundance of taxon *i* across communities; and *I*() is the probability density function of beta distribution. *R*^2^ indicates the fit of the parameter based on nonlinear least-squares fitting.

### 2.7 Data available

Sequencing data are stored on the National Genomics Data Center (NGDC) Sequence Read Archive (PRJCA018972).

## 3. Results

### 3.1 Environmental heterogeneity between urban and natural river areas

A total of 28 urban river and 24 undisturbed pristine natural river surface (0-2 cm) sediment samples were collected (Fig. 1). Samples from all 52 locations covered a wide range of horizontal distance (0.72-376 km). The environmental chemical profiles of river water showed remarkable differences between the urban and natural rivers (Fig. 1, Dataset 1A). The concentration of total nitrogen (TN), nitrite, nitrate, ammonium, total phosphorus (TP), total dissolved phosphorus (TDP), phosphate, volatile phenol, Chlorophyll *a* permanganate index, in urban river were obviously higher than natural river (P<0.01). The total organic carbon (TOC) content in sediment also showed a significant increasing trend from the natural river to urban river (*P*<0.0001) (Fig. 1). Here, the permanganate index is often used as a comprehensive indicator of the degree of water pollution caused by reducible organic and inorganic substances (49).

The sampling locations along the urban river exhibited an average Trophic State Index (TSI) of 53.07, with a range from 41.97 to 69.32 (Dataset 1B), categorizing most of these sites as eutrophic based on a TSI threshold exceeding 50 (50). Conversely, the sampling sites within the natural river had an average TSI of 36.23, with range from 20.84 to 53.95 (Dataset 1B), positioning most sites within the oligotrophic and mesotrophic range (TSI < 50). These differences indicate substantial environmental heterogeneity between the two habitats.

### 3.2 Microbial community abundance, richness, diversity

Microbial abundance ranged from 1.3 × 10^7^ to 1.1 × 10^9^ cells g-1 and 8.0 × 10^6^ to 7.1 × 10^8^ cells g-1 for urban and natural river samples, respectively (Fig. 2A, details in Dataset 2A), and urban river sites exhibited a significantly higher abundance of microorganisms than those from the natural river samples (*P*<0.01). The abundance of microorganisms increased with increasing TSI, indicating that the urban river eutrophication and has a great impact on microbial population (Fig. 2E).

**Figure 2.**
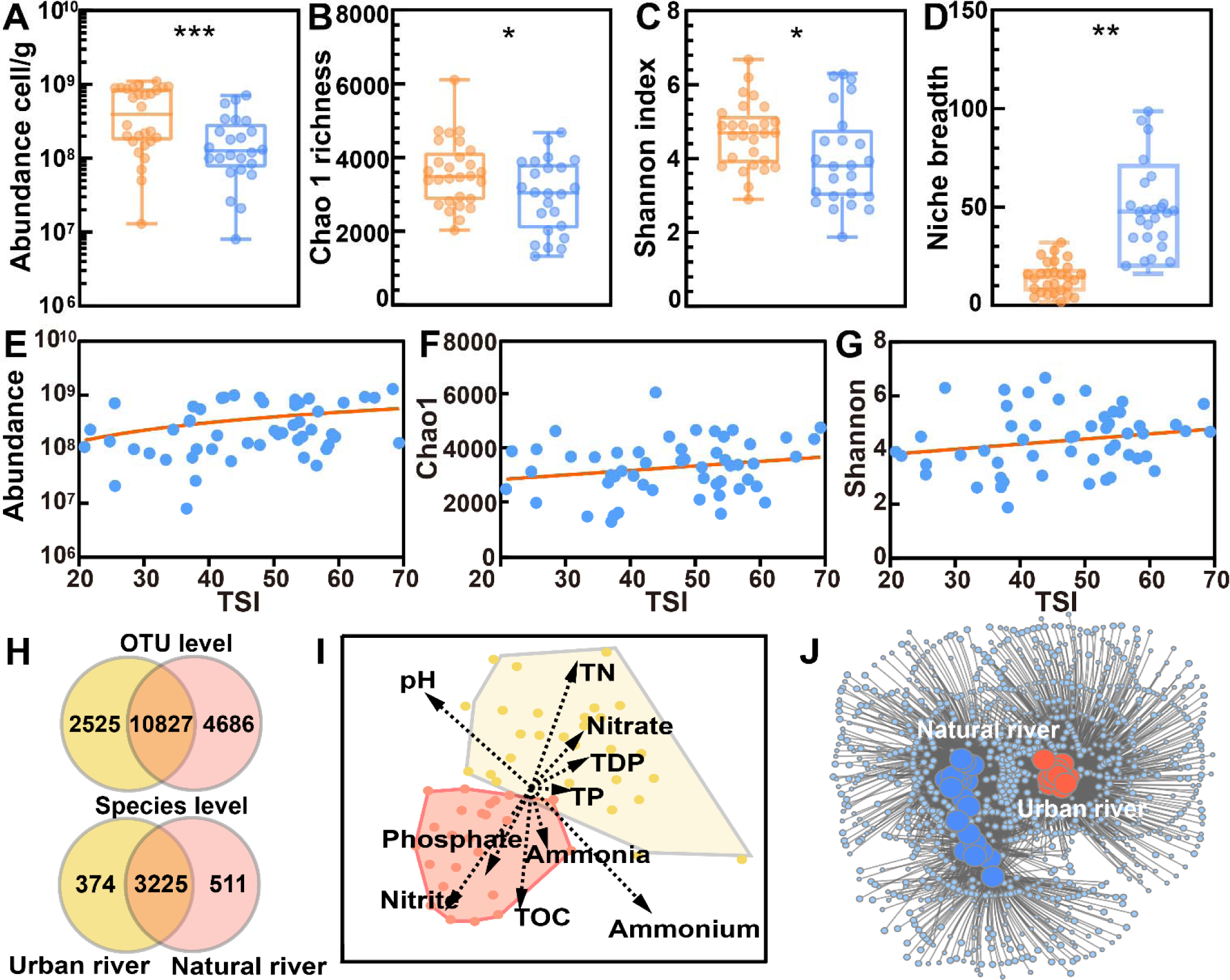
The microbiome in the large river network. (A-D) Microbial abundance, Chao1 richness estimates, Shannon index and Niches breadth were investigated in urban (orange circles) and natural (blue circles) river samples. (E-G) Pairwise relationships between the microbial abundance, Chao 1 richness, Shannon index and TSI for observed communities. (H) Venn analysis. (F) A distance-based redundancy analysis (db-RDA) visualized by two-dimensional (2D) nonmetric multidi-mensional scaling (NMDS) showing microbial community similarity when environmental factors were used as constraining variables and were fitted onto the ordination as arrows. Yellow circles indicate the urban river sampling sites, and red circles indicate the natural river sampling sites, respective. The length of each arrow indicates the multiple partial correlation of the variable to RDA axes and can be interpreted as an indication of that variable’s contribution to the explained community similarity. (I) Co-occurrence networks of the microbial community between urban and natural river samples.

Using universal primers for bacteria and archaea, 3,309,566 high-quality sequences were generated throughout the river network (Dataset 2A, Supporting Information Fig. S1). These high-quality sequences were clustered into 18,038 operational taxonomic units (OTUs, 97% similarity level) (Dataset 2B). A range of 1,628–6,104 chao1 OTUs and 1.89–6.68 Shannon index were observed in the urban samples, and a range of 1,321–5,471 chao1 OTUs and 1.88–6.31 Shannon index were observed in natural river samples (Fig. 2B-C). The richness and diversity were considerably higher in the urban river compared to the natural river samples (*P* < 0.05). The alpha diversity showed predictable patterns along the TSI gradient, and both the Chao 1 and Shannon increased with increasing TSI (Fig. 2F-G). The natural river communities exhibit a wider niche breadth (Fig. 2D), which is indicative of greater metabolic flexibility.

74 microbial phyla and 3,599 species were identified in urban samples, and 80 phyla and 3,736 species were identified in natural samples (Dataset 2C-F). At the phylum level, the urban and natural river samples investigated here shared a high proportion of taxa (Fig. 3A). The ten most prevalent phyla in all of sites collectively constituted 95.48% of the total sequences, matching the classical abundance-range relationship (51, 52). Furthermore, an increase in the diversity and proportion of archaea was also identified under the impact of human activities pollution effluent. In the urban river areas, proportion of archaea exhibited 1.58 times higher than those in natural river, and archaeal Chao1 richness and Shannon diversity exhibited 1.47 and 2.36 times higher than that in natural river, respectively.

**Figure 3.**
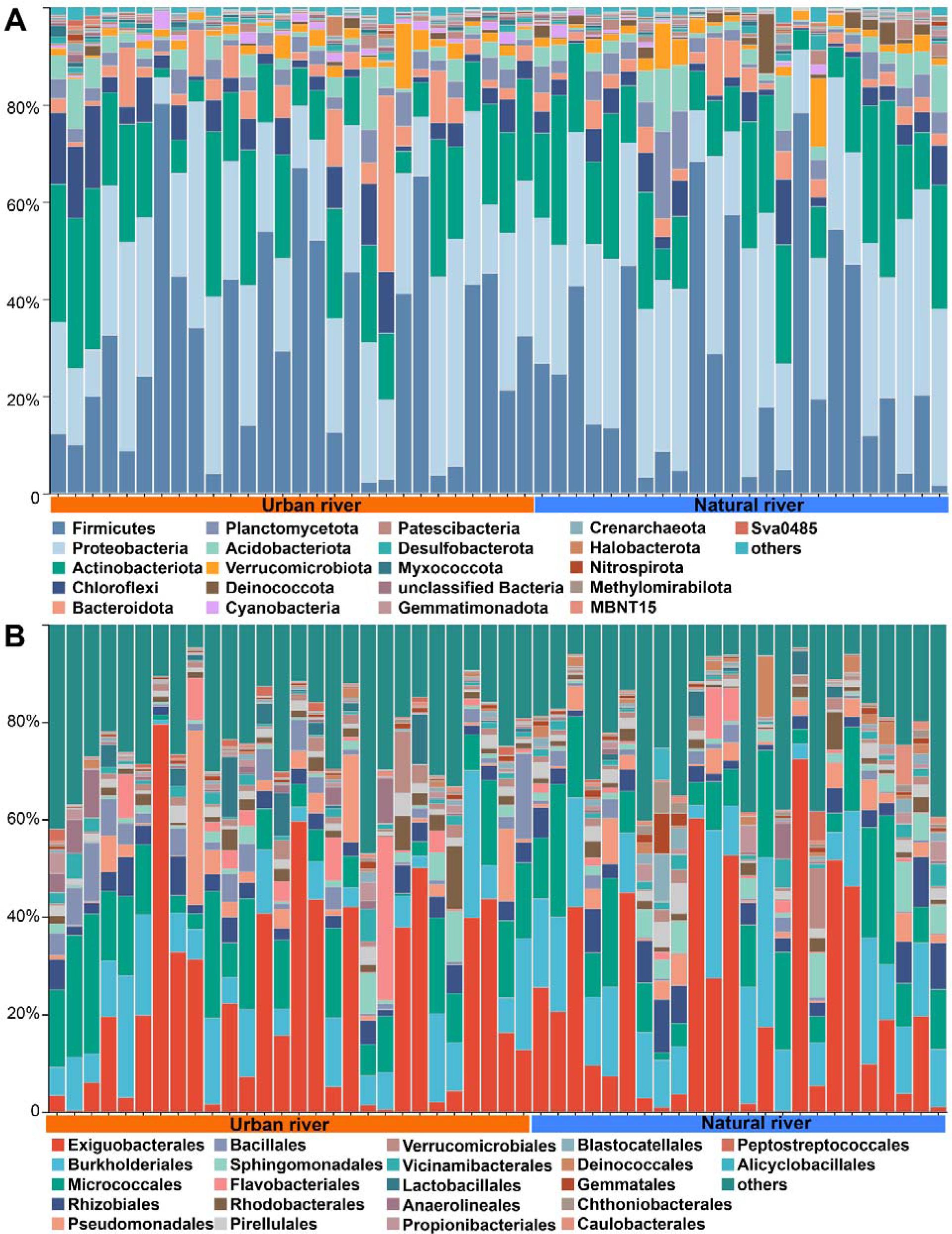
The relative abundance of the phylum (A) and order (B) in urban and natural river sediment samples.

Although the microbial community of these sites exhibited similarities at the phylum level, 38 orders in samples showed significant differences (*P*<0.01) between urban and natural river ecosystems (Supporting Information Fig. S2). For example, Bacillales, Clostridiales, Steroidobacterales were dominant in urban areas, while Erysipelotrichales, Deinococcales, Alicyclobacillales, Pyrinomonadales, Burkholderiales were dominant in the natural river (Dataset 2C, Supporting Information Fig. S2). Further, a heatmap was generated to represent the 50 most prevalent species (Supporting Information Fig. S3). In urban rivers, the dominant taxa included Unclassified Planococcaceae, Unclassified Steroidobacteraceae, Unclassified Anaerolineaceae, and Unclassified Rhizobiales. On the other hand, the unclassified *Acinetobacter*, unclassified Pseudomonas, unclassified Sphingomonadaceae, unclassified *Deinococcus*, *Pseudomonas parafulva* had high abundances in natural rivers (Dataset 2E). At OTU level, the NMDS and network analysis showed that the urban samples harbored significantly dissimilar microbial communities with natural river samples (Fig. 2I-J; PERMANOVA pseudo F = 2.1455, p.adj = 0.024, 999 permutations; ANOSIM R sqr = 0.0597, P = 0.03). Along the trophic gradient, we observed taxonomic homogenization for bacteria and archaea. Such a phenomenon can be supported by the average Bray-Curtis dissimilarity for each trophic group, which showed lower values in highly urbanization river than in natural rivers with lower trophic status.

### 3.3 The key functional microbial communities

Microbial community metabolic predictions guided by the FAPROTAX database (39) revealed distinctions in the metabolic potential related to nitrogen, carbon, and sulfur cycling between urban and natural rivers (Fig. 4, Dataset 3A-D). For nitrogen transformation, the microorganisms involved in ammonia oxidation, nitrite oxidation, nitrogen fixation, nitrification in urban rivers were higher than those in the natural river (ANOVA, *P* < 0.001). High-throughput sequencing showed a notably higher abundance of nitrogen transformation taxa groups (Dataset 2E-F, *P* < 0.01) was observed in urbanized rivers, including nitrogen-fixing bacteria (*Anaeromyxobacte*) (53), nitrifying bacteria (*Nitrosomonas, Nitrospina, Nitrosospira*, *Nitrospira*) and denitrifying bacteria (*Paracoccus, Hyphomicrobium*, and *Gemmobacter*) (54). The db-RDA analysis indicated that nitrogen (often in the form of nitrate and ammonium) contributed significantly to the changes in the urban river microbial community (Fig. 2I).

**Figure 4.**
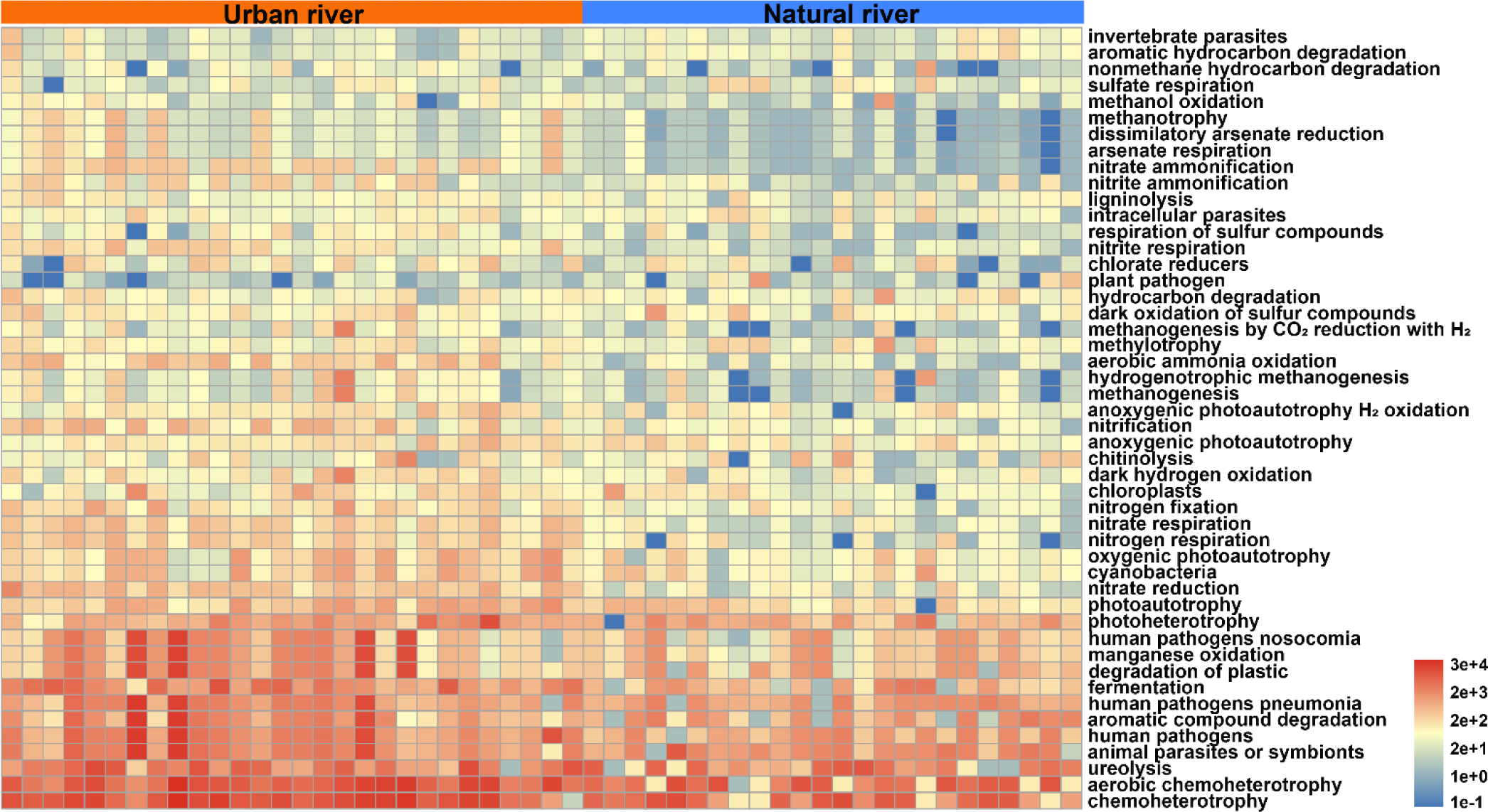
The potentially ecological functions based on 16S rRNA genes of microbial community composition.

In all samples, biodegradation pathways encompassing 21 types of xenobiotic compounds were determined (Dataset 3F-G). The ecological function of xenobiotic biodegradation exhibited a noticeable increasing trend from the natural river to the human-impacted sites. It is understood that the detection of biodegradation genes can act as markers for the presence of xenobiotics and their related metabolites (55, 56). In the urban river locations, a significant presence of aminobenzoate, benzoate, and naphthalene was observed (Dataset 3G), coinciding with a rise in the abundance of bacteria capable of degrading PAHs, such as *Sphingobium*, *Rhodococcus*, *Novosphingobium*, *Pseudomonas*, *Sphingomonas*, *Brevibacterium*, *Arthrobacter*, *Nocardioides,* and *Mycobacterium* were observed in the urban river (Dataset 2E, *P* < 0.01) (57). PAHs primarily originating from long-term anthropogenic pollution sources linked to industrialization and urbanization (58, 59). These PAHs pollutants, characterized by persistence and low solubility, tend to accumulate in river sediments due to their hydrophobic nature and strong adsorption onto sediment particles (5, 58). It suggested the accumulation of PAHs in our collected urban river sediment samples.

Function genes linked to drug and antimicrobial metabolism were observed higher in urban river sediment samples (Dataset 3D). Therefore, three kinds of antibiotic, macrolides, tetracycline and sulfonamide, were selected for antibiotic resistance rates (ARRs) test in our study (Fig. 5A). Tetracyclines and macrolides are the most commonly used antibiotics around the world for both medical and agricultural purposes (60, 61). Sulfonamides, the first antibiotics to be synthesized for widespread use, have experienced a significant rise in resistance among clinical samples (62). The ARRs in urban rivers ranged 11.2% - 29.5 % for azithromycin, 13.0-29.0 % for tetracycline, 11.0-26.9 % for sulfonamide. The ARRs in natural rivers ranged 2.4-18.2% for azithromycin, 4.0-16.0 % for tetracycline, 2.4-16.0 % for sulfonamide. The antibiotic resistant rate in urban samples were higher than those in natural river samples (Fig. 5A). Five antibiotic resistance genes (ARGs) spanning three categories of resistance mechanisms—specifically, two for sulfonamides (*sul1* and *sul2*), two for tetracyclines (*tetO* and *tetW*), and one for macrolides (*ermF*) (Fig. 5B)—were chosen for analysis in all specimens to delineate the variance in ARG concentrations between urban and natural river settings. The ARGs were found in all samples, and Fig. 5B shows the occurrences and quantities of ARGs in the sediments of the urban and natural rivers. The concentrations of ARGs varied greatly, ranging from 1.04 × 10^3^ copies/g (*sul2*, Natural River) to 4.10 × 10^8^ copies/g (*tetO*, urban river). The quantity of ARGs in the urban river ranged from 1.36 × 10^4^ to 1.63 × 10^8^ copies/g for *sul1*, 2.27 × 10^4^ to 3.00 × 10^8^ copies/g for *sul2*, 5.36 × 10^3^ to 4.10 × 10^8^ copies/g for *tetO,* 2.52 × 10^3^ to 2.17 × 10^8^ copies/g for *tetW,* 1.72 × 10^3^ to 2.02 × 10^8^ copies/g for *ermF*. The quantity of ARGs in the natural river ranged from 1.42 × 10^3^ to 9.49 × 10^6^ copies/g for *sul1*, 1.04 × 10^3^ to 1.9 × 10^7^ copies/g for *sul2*, 1.49 × 10^3^ to 2.5 × 106 copies/g for *tetO,* 2.34 × 10^3^ to 9.45 × 106 copies/g for *tetW, 2.12* × 10^3^ to 5.52 × 10^6^ copies/g for *ermF*. The quantity of ARGs abundances in the sediments of the urban river were higher than those in the sediments collected from the natural river.

**Figure 5.**
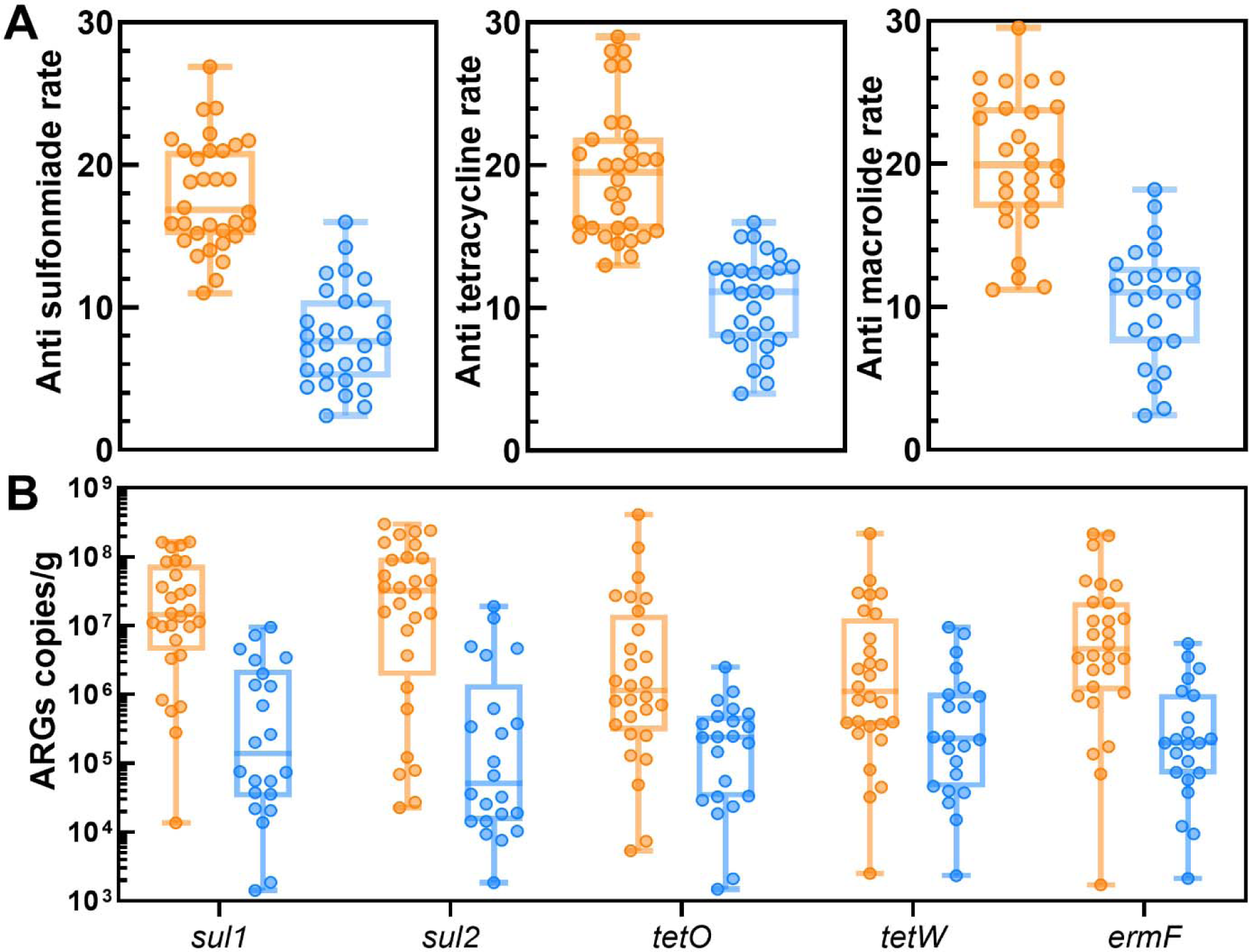
Different of antibiotic resistance rates (A) and antibiotic resistance gene abundance (B). The orange circles indicate the urban river and blue circles indicate the natural river, respectively.

Further, the urban areas displayed a higher abundance of taxa involved in fecal contamination and pathogens compared to the natural river (ANOVA, *P* < 0.05). For example, high abundance of Bacteroidetes and Firmicutes were observed in the urban river (Dataset 2C, *P* <0.01). Firmicutes and Bacteroidetes are two major phyla of bacteria found in the human intestine, and have been frequently found in WWTPs and wastewater-influenced environments (63), which are recognized as good indicators of fecal contamination (40, 64). The elevated abundance of Bacteroidetes and Firmicutes in our collected urban river sediment samples suggested the presence of untreated wastewater from residential areas in these areas, which may contain fecal pollutants. Additionally, according to Pathogen–Host Interaction (PHI) database (65), our study also identified a marked increase in the abundance and diversity of pathogens like *Acinetobacter*, *Arcobacter*, *Exiguobacterium*, *Chryseobacterium*, *Escherichia Shigella* and *Staphylococcus aureus* within urban river (Dataset 2E-F, P < 0.01). These findings highlighted urban non-point source untreated water pollutants have resulted in fecal and pathogenic bacterium contamination in the river network.

### 3.6 Key influencing factors and ecological driving processes

Across the large river network spanning a geographic distance of ∼400 km, the microbial community increased in dissimilarity with increasing spatial distance, showed a distance-decay relationship (DDR) (Fig. 6A, Dataset 2G). However, we observed microbial communities in the natural rivers had a steeper DDR slope between sampled sites than those in the urban rivers (Fig. 6A), indicating the environmental selection has a stronger effect on the urban river. The distance-based redundancy analysis (db-RDA) using environmental parameters as constraining variables, highlighted the potential significance of these chemical parameters in elucidating microbial community compositions (Fig. 2I). Ammonium (r sqr= 0.3551, *P*=0.001), pH (r sqr= 0.2293, *P*=0.005), Nitrite (r sqr=0.2211, *P*=0.003), Nitrate (r sqr=0.188 *P*=0.006), TN (r sqr= 0.17, *P*=0.006), TOC (r sqr= 0.1367, *P*=0.032), phosphate (r sqr= 0.0615, *P*=0.010) were more crucial in shaping the microbial community.

**Figure 6.**
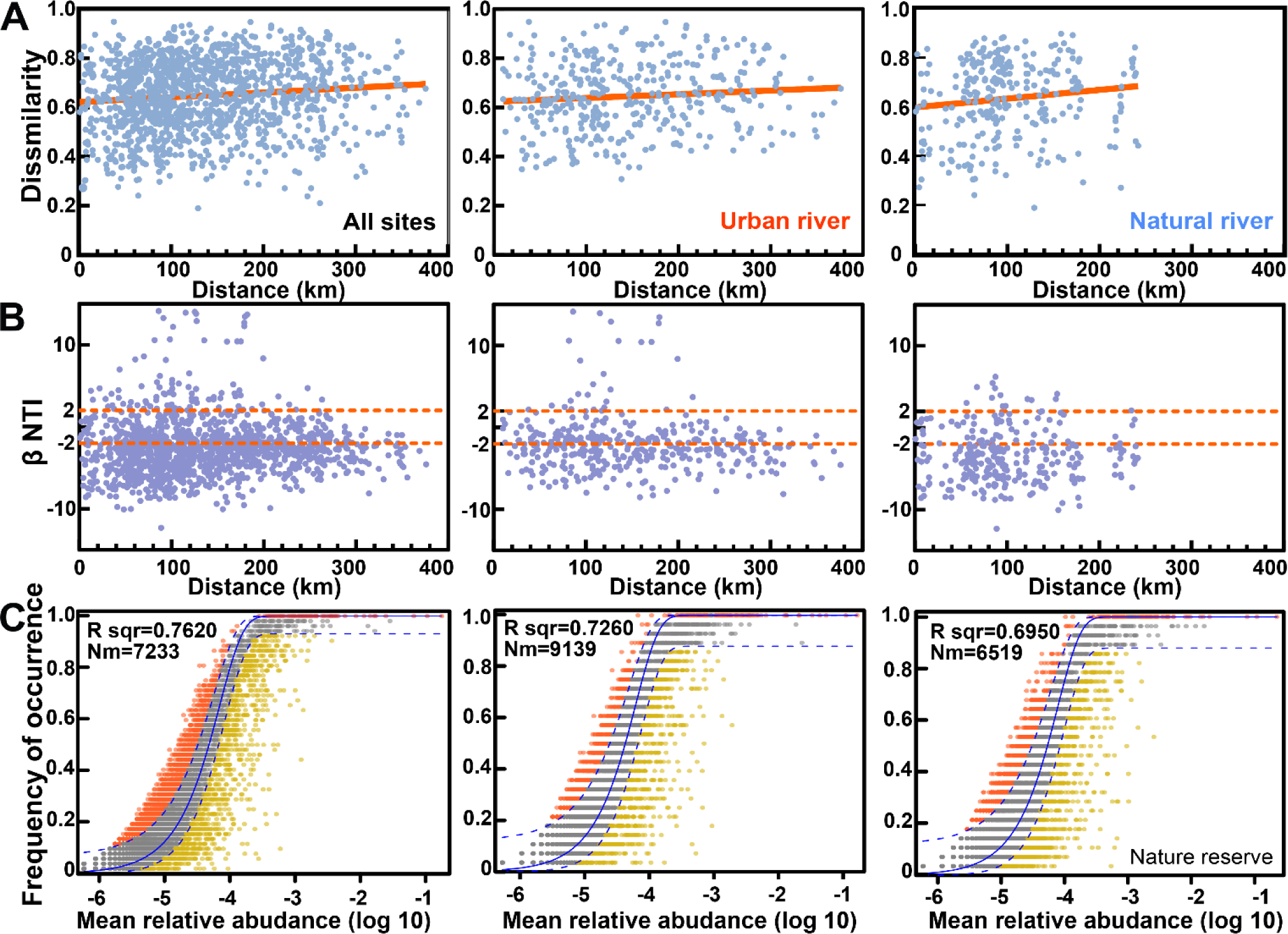
General patterns of microbial diversity in the river network system. (A) Distance-decay curves showing Bray–Curtis dissimilarity against geographic distances between sampling sites. Solid lines denote the ordinary least-squares linear regressions. (B) Scatter plot of βNTI values grouped by spatial scales of the river network. (C) Fit of the neutral community model (NCM) of community assembly. The solid blue lines indicate the best fit to the NCM as in Sloan et al. (2006), and the dashed blue lines represent 95% confidence intervals around the model prediction. OTUs that occur more or less frequently than predicted by the NCM are shown in different colors. Nm indicates the metacommunity size times immigration, R^2^ indicates the fit to this model.

To further assess the underlying factors governed community dynamics in the river network, the null model (66) and neutral community model (NCM) (48) were used to quantify the relative influences of stochastic and deterministic processes in our study. The null model indicated a prevalence of environmental selection processes, particularly in the form of homogeneous selection (|βNTI| < 2, RCbary >+0.95) (Fig. 6B), in governing the urban river community assembly (details in Materials and Methods). Simultaneously, neutral community model (Fig. 6C) showed a gradual increase in the contribution of deterministic process with the increase of human activity and eutrophication, elucidating48.3% and 58.5% of the stochastic process within the natural river and urban river samples, respectively. The NCM and null model further revealed that deterministic process, specifically homogeneous selection, are the primary drivers of bacterial and archaeal community compositions in urban river sediments.

## 4. Discussion

In this study, we explored the bacterial and archaeal communities in urban and natural rivers along the trophic gradient and examined the drivers underlying the observed alpha and beta diversity. Considerable variation exists within the bacterial and archaeal communities in terms of abundance, richness, diversity (Fig. 2A-C) and composition (Fig. 2H-J) between urban and natural rivers, and the urbanization enhance microbial population and biodiversity, accompanied by significant niche differentiation (Fig. 2D). Nonmetric multidimensional scaling (NMDS) (Fig. 2I) and network analysis (Fig. 2J), based on OTUs, demonstrated distinct differences between the microbial communities of urban and natural river ecosystems, indicating that community composition trends became increasingly similar as eutrophication intensified. The impact of human activities on river microbial communities was also evident in the predicted metabolic functions of these communities. This differentiation persisted even when considering the exclusion of “rare biosphere” taxa (< 0.1% total sequence).

We inferred that the observed biotic homogenization caused by urbanization eutrophication result from the habitat heterogeneity and environmental filtering. In highly eutrophic urban river systems, the rapid urbanization tends to decrease the overall habitat heterogeneity. For instance, the high nitrogen/phosphorus loading, high organic carbon and lower pH (Fig. 1), are common consequences of the continuous input of industrial, domestic, and agricultural wastewater, which was caused by a large population of approximately 11.06 million in sampled regions (19). The db-RDA analysis (Fig. 2I) indicated that pH and nitrogen (often in the form of high abundance of nitrate and ammonium) contributed significantly to the changes in the urban river microbial community. pH is widely recognized as a major environmental driver shaping microbial communities, influencing the balance and interaction between niche-related and neutral processes (67). More acidic conditions (Fig. 1) imposed stronger environmental filtering and largely shifted community composition. Meanwhile, excessive nutrition inputs lead to eutrophication (Dataset 1B), a process where the nutrient enrichment promotes the growth of nitrogen-transformation microorganisms and algae (Chlorophyll *a* content in Fig. 1) (68). The proliferation of nitrogen-transformation microorganisms and algae leads to increased organic matter production (TOC content in Fig. 1).

Geng et al. (2022) proposed that eutrophication can induce homogenization by modulating interspecies interactions. For instance, organic carbon generated through phytoplankton photosynthesis serves as a crucial carbon source for river microorganisms. Variations in the organic carbon produced by different phytoplankton species select for distinct bacterial communities with varying capacities for organic matter utilization (25). Eutrophication promotes the accumulation of organic matter derived from algae, altering the sources of organic mass and impacting the structure of river microbial communities (69). Additionally, the “productivity/resources hypothesis” suggests that enhanced availability of resources and/or increased primary productivity can sustain larger populations (70). Therefore, the influx of human activity pollutants into the river created a more uniform and homogeneity environment conditions, allowing for the coexistence of a broader range of microbial species. The increased availability of organic matter and the changed nutrient conditions can profoundly influence the river microbial population, and enhanced the microbial population sizes and diversity in urbanized rivers.

Further, higher ARRs and ARGs were also observed in highly urbanized river areas (Fig. 5A-B). Notably, the data showed that significant variation in ARRs and ARGs between cities, with no significant differences in natural river samples. This feature emphasized the importance of the different levels of human activity contribution based on population sizes, potentially resulting in varied ARRs and ARGs among urban rivers. Zhang et al. (2015) reported that over 68% of total antibiotic usage in China in 2013 was excreted by humans and animals, leading to around 53,800 tons of antibiotics entering the environment (71). Urban water ecosystems have been identified as potential hotspots for the spread of antibiotic resistant bacteria in China (72, 73). Hence, the continuous selective pressure exerted by antibiotic residues, even at sub-inhibitory concentrations, in combination with the diverse and dense microbial communities sustained by nutrient-rich environments, creates favorable conditions for the dissemination and proliferation of antibiotic resistance (42).

Meanwhile, nutrient enrichment often coincides with heavy metal pollution in numerous human-impacted aquatic ecosystems (74). In our study, proportion of the archaea in the urban river areas exhibited 1.58 times higher than those in natural river, and archaeal Chao1 richness and Shannon diversity exhibited 1.47 and 2.36 times higher than that in the natural river, respectively. This finding is similar to previous study revealed that in a high heavy metal contamination zone, archaeal Chao1 richness and Shannon diversity exhibited substantial increases, approximately 1.81 and 1.56 times higher than those in a low heavy metal contamination zone, respectively (10). Krzmarzick et al. (2018) proposed that archaea have demonstrated a notable capacity for enduring harsh environments, leading to the possibility that numerous archaeal species possessing exceptional adaptability to severe anthropogenic pollution could flourish in urban river environments (75). Therefore, habitat heterogeneity plays a crucial role in determining alpha and beta diversity in river microbiomes.

Secondly, this study aimed to investigate the assembly mechanisms of both bacterial and archaeal communities in urban and natural rivers using environmental DNA meta-barcoding technology. River microbial communities exhibited high diversity, encompassing thousands of OTUs across major taxa that inhabit various habitats within the river ecosystem (Fig. 2-3). It is worth noting that the distance-decay relationship (DDR) was observed in urban and natural river samples (Fig. 6A). Deciphering the relative influence of diverse ecological processes in microbial community assembly remain an important query in riverine biogeography (76). The strength of the distance–decay of community similarity is influenced by both stochastic and deterministic processes (27, 77). Among them, some previous studies had found the geographic distances (passive dispersal) and environmental filtering are two major ecological processes impacted on the assembly of microbial communities in connected river ecosystem (78-80). For example, in large scale watersheds like the Yangtze River and Tingjiang River, geographic separation and dispersal limitation have been identified as an important factor in the succession of communities (10, 80, 81).

However, in our study, the microbial communities in natural river locations exhibited a more pronounced DDR slope among sampling sites than those in urban river sites (Fig. 6A). The variation in microbial community composition tends to be less pronounced across urban areas compared to the diversity found across natural river ecosystems. Furthermore, our analyses also show narrower niche breadths were observed for samples in the urban river (Fig. 2D), and a higher level of homogenization in the community composition along the urban river compared to the natural river ecosystems (Fig. 2I-J). The application of the neutral community model (Fig. 6C) revealed that deterministic processes predominantly shaped the urban river microbial communities in the river network. Null model further uncovered the homogeneous selection dominated governs the urban river community assembly (Fig. 6B). This finding further suggests that persistent uniform non-point source pollution such as high nutrients, PAHs, antibiotics pollutions arises from the urbanization created a more uniformity environment in urban river system, exerting robust homogeneous selection on urban river ecosystem. Homogeneous selection is likely a potent mechanism for biotic homogenization, particularly in harsh environments like those affected by eutrophication, where it diminishes the role of stochastic processes in shaping assemblages (25). Therefore, similar urban river environmental condition could promote community fusion, contributing to a more similar community structure (Fig. 2I-J) and function (Fig.4), and indicating that taxonomic and functional homogenization occur simultaneously in the river network.

Our findings, highlighting convergence in the microbiome of urban rivers, align with observed urbanization impacts on communities of bird (82), fish (83), plant (84), invertebrate (85, 86), zooplankton (87), dust samples (88). As outlined by Olden and Poff (2003), functional homogenization occurs when a specific type of environment becomes more uniformly common. Consequently, the proliferation of analogous species and consistent physical environments in heavily altered urban river habitats is likely lead to a rise in functional homogenization.

On the other hand, differences in the local species pool between urban and natural river areas were evident, with shared microorganisms accounting for 55% (Fig. 2H). This further underscore distinct environmental conditions between these study areas result in varied local species pools and species abundance distributions. The urban river habitats promote the establishment of 374 “urban-adaptable” species (Fig. 2H), and these “urban-adaptable” species dominate across urban river sites. Meanwhile, a narrower niche breadth has been found in urban river areas compared to natural river areas (Fig. 2D), suggesting a lesser metabolic plasticity of microorganisms. We hypothesize that urban eutrophication favors species adapted to pollution tolerance, thereby promoting biotic homogenization.

In natural ecosystems, microorganisms are typically categorized as habitat generalists and specialists based on their niche breadth (89). Habitat generalists with broad niche breadth are less competitive under optimal growth conditions but exhibit greater resistance to changing environments, whereas specialists have narrower niches and demonstrate specific environmental fitness (90). We had confirmed that anthropogenic activities decreasing urban local environmental heterogeneity in interconnected river network on a regional scale. This decreased environmental heterogeneity not only deteriorates the habitats of indigenous species that are at a competitive disadvantage but also fosters urban-adaptable species (including the exotic species or range expansion of native specialist species) acclimatizing to urban river conditions (Fig. 2D, H) (91). Hence, the mechanisms of biotic homogenization can be attributed to three aspects: (1) extinction of native generalist species, (2) invasion of widespread nonnative species, and (3) range expansion of native specialist species.

Non-native (exotic) biota and native specialist species becomes increasingly enriched in urban rivers, replacing many natural river locally unique, endemic native species from natural rivers, thereby reducing the unique regional identity of biotas. For example, urbanization-induced biotic homogenization leads to an increase in local biodiversity within urban areas (Fig. 2B-C), yet it contributes to a decrease in regional biodiversity in the urban areas (e.g., 3499 species in the urban areas, 3736 species in natural areas, Fig. 2H). It’s worth noting that global diversity diminishes due to the extinction of unique regional species, which are then absent from the worldwide pool of species (92). Therefore, the biotic homogenization could potentially have detrimental effects on crucial ecosystem functions. The finding serves as a compelling example of biotic homogenization, highlighting how human-induced environmental changes are diminishing the biological diversity that exists between natural ecosystems. This trend is expected to lead to a more uniform biosphere at both regional and global levels (92), consequently diminishing global biodiversity. This contrast between local augmentation and global diminution of biodiversity represents a critical concern for conservation biology, as it could potentially divert public attention from the more pressing issue of worldwide species loss.

## 5. Conclusion

Collectively, our research demonstrated that human population and contamination level serve as the predominant influences shaping urban river microbial communities in the river network. Changed environmental conditions can potentially favor specific bacterial and archaeal taxa, alter the alpha and beta diversity of urban river microbiomes, and contribute to the biotic homogenization of river microbiomes. Biotic homogenization can be attributed to the modulation of generalist/specialist species and the invasion of nonnative species. These changes have cascading effects on ecosystem functioning and overall microbial processes in the river network, potentially profoundly altering riverine ecosystems. Future research is required to assess the extent of biotic homogenization due to human activities across global river ecosystems.

## Acknowledgements

This work was supported by the Youth Program of National Natural Science Foundation of China (No. 32100081), Central Public-Interest Scientific Institution Basal Research Fund, CAFS (NO.14).

## Declaration of Generative AI and AI-assisted technologies in the writing process

Statement: During the preparation of this work the authors used CHATGPT in order to improve readability and language. After using this tool/service, the authors reviewed and edited the content as needed and takes full responsibility for the content of the publication.

## Reference

1. Widder S, Besemer K, Singer GA, Ceola S, Bertuzzo E, Quince C, et al. Fluvial network organization imprints on microbial co-occurrence networks. Proceedings of the National Academy of Sciences. 2014;111(35):12799–804.

2. Beattie RE, Bandla A, Swarup S, Hristova KR. Freshwater sediment microbial communities are not resilient to disturbance from agricultural land runoff. Front Microbiol. 2020;11:539921.

3. Wang Y, Sheng HF, He Y, Wu JY, Jiang YX, Tam NF, et al. Comparison of the levels of bacterial diversity in freshwater, intertidal wetland, and marine sediments by using millions of illumina tags. Appl Environ Microbiol. 2012;78(23):8264–71.

4. Singh R, Singh AP, Kumar S, Giri BS, Kim K-H. Antibiotic resistance in major rivers in the world: A systematic review on occurrence, emergence, and management strategies. Journal of Cleaner Production. 2019;234:1484–505.

5. Patel AB, Shaikh S, Jain KR, Desai C, Madamwar D. Polycyclic Aromatic Hydrocarbons: Sources, Toxicity, and Remediation Approaches. Front Microbiol. 2020;11:562813.

6. Xu N, Hu H, Wang Y, Zhang Z, Zhang Q, Ke M, et al. Geographic patterns of microbial traits of river basins in China. Sci Total Environ. 2023;871:162070.

7. Liu Y, Su B, Mu H, Zhang Y, Chen L, Wu B. Effects of point and nonpoint source pollution on urban rivers: From the perspective of pollutant composition and toxicity. J Hazard Mater. 2023;460:132441.

8. Pruden A, Arabi M, Storteboom HN. Correlation Between Upstream Human Activities and Riverine Antibiotic Resistance Genes. Environ Sci Technol. 2012;46(21):11541–9.

9. Manyi-Loh CE, Mamphweli SN, Meyer EL, Makaka G, Simon M, Okoh AI. An Overview of the Control of Bacterial Pathogens in Cattle Manure. International Journal of Environmental Research and Public Health. 2016;13(9):843.

10. Yang S, Chen Q, Zheng T, Chen Y, Zhao X, He Y, et al. Multiple metal(loid) contamination reshaped the structure and function of soil archaeal community. J Hazard Mater. 2022;436:129186.

11. Delgado-Baquerizo M, Eldridge DJ, Liu Y-R, Sokoya B, Wang J-T, Hu H-W, et al. Global homogenization of the structure and function in the soil microbiome of urban greenspaces. Science Advances. 2021;7(28):eabg5809.

12. Drury B, Rosi-Marshall E, Kelly JJ. Wastewater treatment effluent reduces the abundance and diversity of benthic bacterial communities in urban and suburban rivers. Appl Environ Microbiol. 2013;79(6):1897–905.

13. Pascual-Benito M, Ballesté E, Monleón-Getino T, Urmeneta J, Blanch AR, García-Aljaro C, et al. Impact of treated sewage effluent on the bacterial community composition in an intermittent mediterranean stream. Environ Pollut. 2020;266:115254.

14. Dai T, Su Z, Zeng Y, Bao Y, Zheng Y, Guo H, et al. Wastewater treatment plant effluent discharge decreases bacterial community diversity and network complexity in urbanized coastal sediment. Environ Pollut. 2023;322:121122.

15. Kiedrzyńska E, Kiedrzyński M, Urbaniak M, Magnuszewski A, Skłodowski M, Wyrwicka A, et al. Point sources of nutrient pollution in the lowland river catchment in the context of the Baltic Sea eutrophication. Ecol Eng. 2014;70:337–48.

16. Ding X, Liu L. Long-term effects of anthropogenic factors on nonpoint source pollution in the upper reaches of the Yangtze river. Sustainability. 2019;11(8):2246.

17. EPA. Basic Information about Nonpoint Source (NPS) Pollution United States Environmental Protection Agency 2022 [Available from: https://www.epa.gov/nps/basic-information-about-nonpoint-source-nps-pollution.

18. Wang H, Dong Y, Jiang Y, Zhang N, Liu Y, Lu X, et al. Multiple stressors determine the process of the benthic diatom community assembly and network stability in urban water bodies in Harbin. Sci Total Environ. 2024;913:169536.

19. Cheng M, Duan C. The changing trends of internal migration and urbanization in China: new evidence from the seventh National Population Census. China Population and Development Studies. 2021;5:275–95.

20. Dudley N. Guidelines for applying protected area management categories: Iucn; 2008.

21. Xiao H, Zhang X, Yan M, Zhang L, Wang H, Ma Y, et al. The Temporal-Based Forest Disturbance Monitoring Analysis: A Case Study of Nature Reserves of Hainan Island of China From 1987 to 2020. Frontiers in Environmental Science. 2022;10.

22. Gibbons SM, Jones E, Bearquiver A, Blackwolf F, Roundstone W, Scott N, et al. Human and Environmental Impacts on River Sediment Microbial Communities. PLoS One. 2014;9(5):e97435.

23. Battin TJ, Besemer K, Bengtsson MM, Romani AM, Packmann AI. The ecology and biogeochemistry of stream biofilms. Nat Rev Microbiol. 2016;14(4):251–63.

24. Xie Y, Wang J, Wu Y, Ren C, Song C, Yang J, et al. Using in situ bacterial communities to monitor contaminants in river sediments. Environ Pollut. 2016;212:348–57.

25. Geng M, Zhang W, Hu T, Wang R, Cheng X, Wang J. Eutrophication causes microbial community homogenization via modulating generalist species. Water Res. 2022;210:118003.

26. Carlson RE. A trophic state index for lakes 1. Limnol Oceanogr. 1977;22(2):361–9.

27. Li H, Zhou H, Yang S, Dai X. Stochastic and Deterministic Assembly Processes in Seamount Microbial Communities. Applied and environmental microbiology. 2023;89(7).

28. Eickhorst T, Tippkötter R. Improved detection of soil microorganisms using fluorescence in situ hybridization (FISH) and catalyzed reporter deposition (CARD-FISH). Soil Biol Biochem. 2008;40(7):1883–91.

29. Hill JE, Town JR, Hemmingsen SM. Improved template representation in cpn60 polymerase chain reaction (PCR) product libraries generated from complex templates by application of a specific mixture of PCR primers. Environ Microbiol. 2006;8(4):741–6.

30. Hoshino T, Doi H, Uramoto GI, Wormer L, Adhikari RR, Xiao N, et al. Global diversity of microbial communities in marine sediment. Proc Natl Acad Sci U S A. 2020;117(44):27587–97.

31. Li H, Yang Q, Zhou H. Niche differentiation of sulfate-and iron-dependent anaerobic methane oxidation and methylotrophic methanogenesis in deep sea methane seeps. Front Microbiol. 2020;11:1409.

32. Schloss PD, Westcott SL, Ryabin T, Hall JR, Hartmann M, Hollister EB, et al. Introducing mothur: open-source, platform-independent, community-supported software for describing and comparing microbial communities. Appl Environ Microbiol. 2009;75(23):7537–41.

33. Levins R. Evolution in changing environments: some theoretical explorations: Princeton University Press; 1968.

34. Pandit SN, Kolasa J, Cottenie K. Contrasts between habitat generalists and specialists: an empirical extension to the basic metacommunity framework. Ecology. 2009;90(8):2253–62.

35. Jiao S, Yang Y, Xu Y, Zhang J, Lu Y. Balance between community assembly processes mediates species coexistence in agricultural soil microbiomes across eastern China. The ISME Journal. 2020;14(1):202–16.

36. Salazar G. EcolUtils: Utilities for community ecology analysis. R package. 2020.

37. Douglas GM, Maffei VJ, Zaneveld JR, Yurgel SN, Brown JR, Taylor CM, et al. PICRUSt2 for prediction of metagenome functions. Nat Biotechnol. 2020;38(6):685–8.

38. Langille MG, Zaneveld J, Caporaso JG, McDonald D, Knights D, Reyes JA, et al. Predictive functional profiling of microbial communities using 16S rRNA marker gene sequences. Nat Biotechnol. 2013;31(9):814–21.

39. Louca S, Parfrey LW, Doebeli M. Decoupling function and taxonomy in the global ocean microbiome. Science. 2016;353(6305):1272-7.

40. Wang L, Zhang J, Li H, Yang H, Peng C, Peng Z, et al. Shift in the microbial community composition of surface water and sediment along an urban river. Sci Total Environ. 2018;627:600–12.

41. Berendonk TU, Manaia CM, Merlin C, Fatta-Kassinos D, Cytryn E, Walsh F, et al. Tackling antibiotic resistance: the environmental framework. Nat Rev Microbiol. 2015;13(5):310–7.

42. Rizzo L, Manaia C, Merlin C, Schwartz T, Dagot C, Ploy MC, et al. Urban wastewater treatment plants as hotspots for antibiotic resistant bacteria and genes spread into the environment: A review. Sci Total Environ. 2013;447:345–60.

43. Munir M, Wong K, Xagoraraki I. Release of antibiotic resistant bacteria and genes in the effluent and biosolids of five wastewater utilities in Michigan. Water Res. 2011;45(2):681–93.

44. Gao P, Mao D, Luo Y, Wang L, Xu B, Xu L. Occurrence of sulfonamide and tetracycline-resistant bacteria and resistance genes in aquaculture environment. Water Res. 2012;46(7):2355–64.

45. Milaković M, Križanović S, Petrić I, Šimatović A, González-Plaza JJ, Gužvinec M, et al. Characterization of macrolide resistance in bacteria isolated from macrolide-polluted and unpolluted river sediments and clinical sources in Croatia. Sci Total Environ. 2020;749:142357.

46. Brooks J, Maxwell S, Rensing C, Gerba C, Pepper I. Occurrence of antibiotic-resistant bacteria and endotoxin associated with the land application of biosolids. Can J Microbiol. 2007;53(5):616–22.

47. Li H, Yang Q, Li J, Gao H, Li P, Zhou H. The impact of temperature on microbial diversity and AOA activity in the Tengchong Geothermal Field, China. Sci Rep. 2015;5:17056.

48. Sloan WT, Lunn M, Woodcock S, Head IM, Nee S, Curtis TP. Quantifying the roles of immigration and chance in shaping prokaryote community structure. Environ Microbiol. 2006;8(4):732–40.

49. Xie W-Q, Gong Y-X. Measurement of permanganate index in environmental water via indirect phase-conversion strategy. Journal of Chromatography A. 2024;1728:464987.

50. Zhang J, Zhang B, Liu Y, Guo Y, Shi P, Wei G. Distinct large-scale biogeographic patterns of fungal communities in bulk soil and soybean rhizosphere in China. Sci Total Environ. 2018;644:791–800.

51. Ruff SE, Biddle JF, Teske AP, Knittel K, Boetius A, Ramette A. Global dispersion and local diversification of the methane seep microbiome. Proc Natl Acad Sci U S A 2015;112(13):4015–20.

52. Amend AS, Oliver TA, Amaral-Zettler LA, Boetius A, Fuhrman JA, Horner-Devine MC, et al. Macroecological patterns of marine bacteria on a global scale. J Biogeogr. 2013;40(4):800–11.

53. Masuda Y, Yamanaka H, Xu Z-X, Shiratori Y, Aono T, Amachi S, et al. Diazotrophic anaeromyxobacter isolates from soils. Applied and environmental microbiology. 2020;86(16):e00956–20.

54. Lücker S, Wagner M, Maixner F, Pelletier E, Koch H, Vacherie B, et al. A Nitrospira metagenome illuminates the physiology and evolution of globally important nitrite-oxidizing bacteria. Proc Natl Acad Sci U S A. 2010;107(30):13479–84.

55. Pagé AP, Yergeau É, Greer CW. Salix purpurea Stimulates the Expression of Specific Bacterial Xenobiotic Degradation Genes in a Soil Contaminated with Hydrocarbons. PLoS One. 2015;10(7):e0132062.

56. Kansole MMR, Lin T-F. Microcystin-LR Biodegradation by Bacillus sp.: Reaction Rates and Possible Genes Involved in the Degradation. Water. 2016;8(11):508.

57. Ghosal D, Ghosh S, Dutta TK, Ahn Y. Corrigendum: Current State of Knowledge in Microbial Degradation of Polycyclic Aromatic Hydrocarbons (PAHs): A Review. Front Microbiol. 2016;7:1837.

58. Kuppusamy S, Thavamani P, Venkateswarlu K, Lee YB, Naidu R, Megharaj M. Remediation approaches for polycyclic aromatic hydrocarbons (PAHs) contaminated soils: Technological constraints, emerging trends and future directions. Chemosphere. 2017;168:944–68.

59. Mojiri A, Zhou JL, Ohashi A, Ozaki N, Kindaichi T. Comprehensive review of polycyclic aromatic hydrocarbons in water sources, their effects and treatments. Sci Total Environ. 2019;696:133971.

60. Van Boeckel TP, Gandra S, Ashok A, Caudron Q, Grenfell BT, Levin SA, et al. Global antibiotic consumption 2000 to 2010: an analysis of national pharmaceutical sales data. The Lancet infectious diseases. 2014;14(8):742-50.

61. WHO. Critically important antimicrobials for human medicine. 6th Revision 2018: World Health Organization. 2019.

62. Sköld O. Sulfonamide resistance: mechanisms and trends. Drug resistance updates. 2000;3(3):155–60.

63. Saarenheimo J, Aalto SL, Rissanen AJ, Tiirola M. Microbial Community Response on Wastewater Discharge in Boreal Lake Sediments. Front Microbiol. 2017;8:750.

64. Wéry N, Lhoutellier C, Ducray F, Delgenès J-P, Godon J-J. Behaviour of pathogenic and indicator bacteria during urban wastewater treatment and sludge composting, as revealed by quantitative PCR. Water Res. 2008;42(1):53–62.

65. Urban M, Cuzick A, Seager J, Wood V, Rutherford K, Venkatesh SY, et al. PHI-base: the pathogen–host interactions database. Nucleic Acids Res. 2020;48(D1):D613–D20.

66. Stegen JC, Lin X, Konopka AE, Fredrickson JK. Stochastic and deterministic assembly processes in subsurface microbial communities. ISME J. 2012;6(9):1653–64.

67. Ren L, Jeppesen E, He D, Wang J, Liboriussen L, Xing P, et al. pH Influences the Importance of Niche-Related and Neutral Processes in Lacustrine Bacterioplankton Assembly. Applied and environmental microbiology. 2015;81(9):3104–14.

68. Kuypers MMM, Marchant HK, Kartal B. The microbial nitrogen-cycling network. Nat Rev Microbiol. 2018;16(5):263–76.

69. Han X, Schubert CJ, Fiskal A, Dubois N, Lever MA. Eutrophication as a driver of microbial community structure in lake sediments. Environ Microbiol. 2020;22(8):3446–62.

70. Ibarbalz FM, Henry N, Brandao MC, Martini S, Busseni G, Byrne H, et al. Global Trends in Marine Plankton Diversity across Kingdoms of Life. Cell. 2019;179(5):1084–97 e21.

71. Zhang Q-Q, Ying G-G, Pan C-G, Liu Y-S, Zhao J-L. Comprehensive Evaluation of Antibiotics Emission and Fate in the River Basins of China: Source Analysis, Multimedia Modeling, and Linkage to Bacterial Resistance. Environ Sci Technol. 2015;49(11):6772–82.

72. Hou L, Zhang L, Li F, Huang S, Yang J, Ma C, et al. Urban ponds as hotspots of antibiotic resistome in the urban environment. J Hazard Mater. 2021;403:124008.

73. Reddy S, Kaur K, Barathe P, Shriram V, Govarthanan M, Kumar V. Antimicrobial resistance in urban river ecosystems. Microbiol Res. 2022;263:127135.

74. Jaiswal D, Pandey J. An ecological response index for simultaneous prediction of eutrophication and metal pollution in large rivers. Water Res. 2019;161:423–38.

75. Krzmarzick MJ, Taylor DK, Fu X, McCutchan AL. Diversity and Niche of Archaea in Bioremediation. Archaea. 2018;2018:3194108.

76. Martiny JB, Eisen JA, Penn K, Allison SD, Horner-Devine MC. Drivers of bacterial beta-diversity depend on spatial scale. Proc Natl Acad Sci U S A. 2011;108(19):7850–4.

77. Stegen JC, Lin X, Fredrickson JK, Chen X, Kennedy DW, Murray CJ, et al. Quantifying community assembly processes and identifying features that impose them. ISME J. 2013;7(11):2069–79.

78. Dini-Andreote F, Stegen JC, van Elsas JD, Salles JF. Disentangling mechanisms that mediate the balance between stochastic and deterministic processes in microbial succession. Proc Natl Acad Sci U S A. 2015;112(11):1326–32.

79. Jia X, Dini-Andreote F, Falcão Salles J. Community assembly processes of the microbial rare biosphere. Trends Microbiol. 2018;26(9):738–47.

80. Chen W, Ren K, Isabwe A, Chen H, Liu M, Yang J. Stochastic processes shape microeukaryotic community assembly in a subtropical river across wet and dry seasons. Microbiome. 2019;7(1):138.

81. Gao Y, Zhang W, Li Y, Wu H, Yang N, Hui C. Dams shift microbial community assembly and imprint nitrogen transformation along the Yangtze River. Water Res. 2021;189:116579.

82. Murthy AC, Fristoe TS, Burger JR. Homogenizing effects of cities on North American winter bird diversity. Ecosphere. 2016;7(1):e01216.

83. Menezes RF, Borchsenius F, Svenning JC, Davidson TA, Søndergaard M, Lauridsen TL, et al. Homogenization of fish assemblages in different lake depth strata at local and regional scales. Freshwater Biology. 2015;60(4):745–57.

84. Salgado J, Sayer CD, Brooks SJ, Davidson TA, Goldsmith B, Patmore IR, et al. Eutrophication homogenizes shallow lake macrophyte assemblages over space and time. Ecosphere. 2018;9(9):e02406.

85. McKinney ML. Effects of urbanization on species richness: a review of plants and animals. Urban ecosystems. 2008;11:161–76.

86. Donohue I, Jackson AL, Pusch MT, Irvine K. Nutrient enrichment homogenizes lake benthic assemblages at local and regional scales. Ecology. 2009;90(12):3470–7.

87. Liu P, Xu S, Lin J, Li H, Lin Q, Han B-P. Urbanization increases biotic homogenization of zooplankton communities in tropical reservoirs. Ecological Indicators. 2020;110:105899.

88. Barberán A, Ladau J, Leff JW, Pollard KS, Menninger HL, Dunn RR, et al. Continental-scale distributions of dust-associated bacteria and fungi. Proc Natl Acad Sci U S A. 2015;112(18):5756–61.

89. Sun J, Zhou H, Cheng H, Chen Z, Wang Y. Distinct strategies of the habitat generalists and specialists in the Arctic sediments: Assembly processes, co-occurrence patterns, and environmental implications. Mar Pollut Bull. 2024;205:116603.

90. Wang L, Han M, Li X, Yu B, Wang H, Ginawi A, et al. Mechanisms of niche-neutrality balancing can drive the assembling of microbial community. Mol Ecol. 2021;30(6):1492–504.

91. Kowarik I. Novel urban ecosystems, biodiversity, and conservation. Environ Pollut. 2011;159(8):1974–83.

92. McKinney ML. Urbanization as a major cause of biotic homogenization. Biol Conserv. 2006;127(3):247–60.

